# Complement contributes to hyperactive behavior in the 16p11.2 hemideletion mouse model

**DOI:** 10.1101/2025.08.21.671537

**Authors:** Benjamin A. Kelvington, Jaekyoon Kim, Regan Fair, Marie E. Gaine, Ted Abel

**Author notes:** to whom correspondence should be addressed, Address: 169 Newton Road, Iowa City, Iowa 52242. authors contributed equally (Benjamin A. Kelvington, Jaekyoon Kim).

## Abstract

The complement system is a major component of the innate immune system and plays an important role in immune surveillance. Recent research has demonstrated that the complement system also plays pivotal roles in brain development, and dysregulation of complement is involved in neurodegenerative and neuropsychiatric disorders. However, the mechanisms by which the complement system contributes to neurodevelopmental disorders (NDDs) remain poorly understood. In this study, we find that the expression of a central regulator of the complement cascade, complement component 3 (C3), is upregulated in the striatum of mice modeling the 16p11.2 hemideletion (16p11.2 del). 16p11.2 del is among the most common copy number variations associated with NDDs including attention deficit hyperactivity disorder (ADHD), autism spectrum disorder (ASD), and intellectual disability (ID). Pharmacological inhibition of the C3a receptor alleviates hyperactivity in 16p11.2 del mice, suggesting that elevated complement contributes to NDD-relevant behavioral changes. Due to the pro-inflammatory actions of the C3a receptor, we assess the cytokine environment in the striatum, a key neural substrate for locomotor behavior, and find that several inflammatory factors are upregulated in 16p11.2 del mice. Collectively, these results indicate that increased expression of the complement system, especially C3, mediates hyperactive behavior and is associated with a pro-inflammatory environment in the striatum of 16p11.2 del mice. Our results suggest that inhibition of an overactive complement system may be an effective strategy to ameliorate NDD symptoms resulting from 16p11.2 hemideletion including those associated with ADHD.

## 1. Introduction

Neurodevelopmental disorders (NDDs) encompass a broad spectrum of conditions that emerge early in development and produce pervasive impairments in functioning (1,2). NDDs including Autism Spectrum Disorder (ASD), Attention-Deficit/Hyperactivity Disorder (ADHD), and Intellectual Disability (ID) are highly prevalent and are each estimated to affect between 1.4% and 8.5% of children (3). There is a critical lack of therapeutic interventions for NDDs as the molecular etiologies underlying these conditions remain unknown. Rare genetic variations associated with NDDs create the opportunity to study these molecular etiologies in animal models. Deletion of the 16p11.2 region (16p11.2 del) is a copy number variation that results in a high penetrance of NDDs including ASD (24%), ADHD (29%), and ID (30%) (4). The 7qF3 region of the mouse genome is syntenic to the human 16p11.2 region allowing for the generation of a genetically valid model of 16p11.2 del in mice (5). These mice display numerous behaviors relevant to NDDs including hyperactivity, learning deficits, motor delays, and disrupted sleep that mirror those observed in humans with 16p11.2 del (5–8). Thus, 16p11.2 del mice represent an important model with translational potential for revealing mechanisms amenable to therapeutic targeting in individuals with 16p11.2 del.

Immune activation is a promising mechanism implicated in the etiology of NDDs (9,10), although the molecular underpinnings remain undefined. Genetic variation and expression changes in immune genes have been documented in NDDs including ASD (11). Furthermore, the study of post-mortem brain tissue of individuals who had ASD reveals activation of microglia, the resident immune cells of the central nervous system (CNS) (12,13). Allergic diseases, characterized by immune overactivation, are also associated with increased risk for an ADHD diagnosis and increased ADHD symptom severity (14). The complement system, a family of well-studied proteins that function as part of the innate immune system, is a promising candidate to explain these neuro-immune associations (15). Recent work has begun to uncover the role of the complement system in the CNS, including how its dysregulation contributes to neurodegenerative and neuropsychiatric disorders (15–18). Remarkably, genes encoding components of the complement system, C1q and C3, are hypomethylated and overexpressed in the prefrontal cortex of individuals with ASD (19). Despite this evidence, the role of complement in animal models for the study of NDDs, including 16p11.2 del, has not been explored. Understanding how neuroimmune mechanisms such as complement activation contribute to NDD symptomatology has the potential to identify targets for the development of novel therapeutic approaches.

The striatum and cortico-striatal circuits are hypothesized to play a critical role in NDDs. The striatum mediates behaviors, such as action initiation, action inhibition, goal-directed learning, habit learning, and cognitive flexibility, that when disrupted can produce behaviors characteristic of NDDs such as impulsivity, hyperactivity, developmental motor delays, and cognitive inflexibility (20–22). Crucially, disruption of cortico-striatal circuitry is a hallmark of both mice modeling 16p11.2 del and humans carrying the deletion (6,23). Prior research reveals that mice modeling 16p11.2 del exhibit robust hyperactivity, a behavior linked to disruptions in striatal function, although the molecular underpinnings of this hyperactivity are unknown (5–7,24,25). The complement system has recently been shown to regulate cortico-striatal synapses in a mouse model for Huntington’s Disease (HD) and is upregulated in post-mortem tissue of human HD samples (26–28). However, the mechanistic role of complement within cortico-striatal circuits in the context of NDDs remains undefined.

Here, we investigate the role of the complement system in NDD-relevant behavior in the 16p11.2 del mouse model. We find that complement is upregulated selectively in the 16p11.2 del striatum, and pharmacological inhibition of the complement system ameliorates the hyperactivity of 16p11.2 del mice. We also identify a pro-inflammatory environment within the 16p11.2 del striatum marked by cytokine production and microglial activation. These findings provide insight into the cellular and molecular mechanisms mediating NDD-relevant behavior in 16p11.2 del mice and pave the way for future therapeutic strategies targeting the neuroimmune mechanisms underlying NDDs.

## 2. Material and methods

### Animals

16p11.2 del mice were produced by breeding male B6129S-Del(7Slx1bSept1)4Aam/J mice purchased from The Jackson Laboratory (Stock #013128) with female B6129SF1/J hybrid mice (Stock #101043). Experimental data was collected from 16p11.2 del animals and wildtype (WT) littermates aged between 10 and 16 weeks. Data from both males and females was combined unless otherwise specified. Mice were housed in a room with a 12-hour light/dark cycle and were allowed *ad libitum* access to food and water. All procedures were approved by the University of Iowa Institutional Animal Care and Use Committee and followed policies set forth by the National Institutes of Health Guide for the Care and Use of Laboratory Animals.

### Home Cage Activity Monitoring

To measure locomotor activity, an infrared beam-break system (Opto M3, Columbus Instruments, Columbus, OH) was employed as previously described (7). Mice were habituated to the monitoring chambers for one week prior to data collection.

### Pharmacology

Beginning in the second week of activity monitoring, animals were injected with vehicle at the beginning of the dark cycle (ZT12). Vehicle injections were delivered at 5 ml/kg, equivalent to the dose used in the subsequent drug delivery schedule. For the C3aR antagonist study 16% DMSO in 0.9% saline was used as a vehicle. 0.9% saline was used as a vehicle for the minocycline study. Beginning on the third week of monitoring, injections were given daily at the onset of the dark cycle (ZT12). For the C3aR antagonist study SB290157 (Sigma-Aldrich, SML1192) was delivered in 2 mg/ml solution for a 10 mg/kg final dose. Minocycline (Tocris, 3268) was delivered in a 10 mg/ml solution to achieve a dose of 50 mg/kg.

### Quantitative Polymerase Chain Reaction (qPCR)

Mice were sacrificed via cervical dislocation and rapidly microdissected. Coronal tissue sections (2 mm) encompassing the striatum were removed, and cortical and striatal samples were taken from this slice. Striatal sections included both dorsal and ventral striatum. Striatum and cortical fractions were submersed in Invitrogen™ RNAlater™ Stabilization Solution (AM7020). RNA extraction was completed using the RNeasy Plus Mini Kit (Qiagen, 74134). Tissue homogenization was performed using TissueRuptor II (Qiagen, 9002755) in RLT lysis buffer. cDNA was prepared using the BioRad iScript cDNA Synthesis Kit (1708890). cDNA reaction was performed using BioRad T100 ThermoCycler according to kit instructions. cDNA was diluted 1:10. Reaction used Applied Biosystems™ Fast SYBR™ Green Master Mix (4385610) and oligonucleotide primers (obtained from Integrated DNA Technologies) with sequences listed in Supplementary Table 1. qPCR was performed using Applied Biosystems QuantStudio 7 Flex instrument and QuantStudio Real-Time PCR software. Data were analyzed using the ΔΔCT method with *Tubulin* and *Gapdh* used as reference genes.

### Enzyme-linked Immunosorbent Assay ELISA

ELISA was performed following manufacturer’s instructions from the following kits: Mouse C3 SimpleStep ELISA Kit (Abcam ab263884) and Mouse C1q ELISA Kit (MyBioSource MBS2702391). Samples were rapidly microdissected from the cortex and striatum, flash frozen, and remained on dry ice until immediately prior to the addition of extraction buffer. Serum samples were also collected. Bradford Reagent was used for protein quantification (Bio-Rad Protein Assay Dye Reagent Concentrate, 5000006). BSA standards ranging from 0-8 µg were used to generate a standard curve, and absorbance at 595 nm was read using a Fisher Scientific AccuSkan GO plate reader.

### Inflammatory Cytokine Array

Assay was performed using the Mouse Inflammation Antibody Array - Membrane (ab133999) and following the protocol recommended by the manufacturer. During protein extraction, 1mL of lysis buffer was supplemented with Halt Protease and Phosphatase Inhibitor Cocktail (100X) (Thermo #1861281). Samples were rapidly microdissected from the striatum and flash frozen then remained on dry ice until immediately prior to lysis buffer addition. Bradford Reagent was used for protein quantification (Bio-Rad Protein Assay Dye Reagent Concentrate, 5000006). BSA standards ranging from 0-8 µg were used to generate a standard curve, and absorbance at 595 nm was read using a Fisher Scientific AccuSkan GO plate reader.

### Immunohistochemistry and Microglia Morphology Analysis

Animals were perfused with 15mL each of PBS and 4% PFA and collected in 4% PFA fixative. Fixed brains were transitioned to 30% sucrose solution (in PBS) following overnight fixation. 40 µm sections of brain tissue were generated using a cryostat (Leica CM3050 S #7501-07-2019) and placed in PBS with 0.1% sodium azide solution. The tissues were washed 3 times in PBS for 5 minutes, then placed in a blocking solution of 3% normal goat serum and 0.2% Triton for one hour. Tissues were then incubated overnight with 1:1000 rabbit anti-Iba1 (Fujifilm Wako, 019-19741) in blocking solution. Other primary antibodies used were mouse anti-C3aR (Hycult Biotech, HM1123), mouse anti-C3 (Hycult Biotech, HM1045), and rabbit anti-S100β (Abcam, ab41548). Sections were then washed 3 times with PBS for 5 minutes and incubated with 1:500 goat anti-rabbit, AlexaFluor 555 (Invitrogen #A21428) in blocking solution for 2 hours. Additional secondary antibodies used were donkey anti-mouse, AlexaFluor 555 (Invitrogen #A32773) and goat anti-rabbit, AlexaFluor 488 (Invitrogen #A32790). The sections were then washed 3 times with PBS for 5 minutes. The sections were mounted on Superfrost Plus charged slides (Fisher Scientific 12-550-15) and allowed to dry, then coverslipped with Prolong Diamond Antifade mountant with DAPI (Thermo Fisher P36971). Slides were stored in a light-protected slide box at 4°C. Slides were imaged using confocal microscopy (Leica DM6 #5210000769). To capture the maximum projections of microglia, a z-stack of 20 µm was used with 40 steps. The images were taken on 20x magnification with 1.5x zoom. A condensed 2-dimensional image of the z-stack was taken by selecting “maximum projection” in the “Processing” tab of the LAS X software. The MicrogliaMorphology ImageJ macro and R package were used to perform image analysis as described (29). The video tutorial for the MicrogliaMorphology ImageJ macro was followed (video can be found at https://github.com/ciernialab/MicrogliaMorphology). Base code was acquired from https://github.com/ciernialab/MicrogliaMorphologyR.

### Data Analysis

Data visualization and statistics were performed with GraphPad Prism v10.5.0 (La Jolla, CA). Data was analyzed using student’s t-test, and paired t-test, two-way ANOVA with Fisher’s LSD *post-hoc* test as appropriate. For graphs containing data from both male and female animals, closed data points represent data obtained from male animals, and open data points indicate data obtained from female animals.

## 3. Results

### 3.1 The complement system is activated in the striatum of 16p11.2 del mice

To assess the impact of 16p11.2 del on the complement system, we perform qPCR on brain tissue from the striatum and cortex of 16p11.2 del mice and WT littermates. mRNA levels of complement components *C3ar1*, *C3*, *C1qa*, and *C1qb* are increased in the striatum of 16p11.2 del mice (Figures 1A-D). Notably, the level of complement transcripts is unchanged in the cortex (Supplementary Figures 1A-D) suggesting that complement is selectively upregulated in the striatum of 16p11.2 del mice. Furthermore, although sex-biased phenotypes relating to the striatum have been observed in 16p11.2 del mice, complement mRNA expression is not influenced by biological sex (Supplementary Figures 2A-H). *Mapk3*, a gene within the 16p11.2 region, was expressed at roughly half of WT level as expected (Figure 1E). To confirm the activation of the complement system, we perform enzyme-linked immunosorbent assays (ELISAs) to assess protein levels of C1q and C3 in 16p11.2 del animals. Consistent with our qPCR results, we observe the upregulation of complement components C3 and C1q in the striatum (Figures 1F, G). We do not observe changes in complement in either the cortex or in serum (Supplementary Figures 1E-H) suggesting that increased complement is locally produced selectively in the striatum and not the result of systemic complement activation. Together, these data reveal that complement is upregulated in the striatum of 16p11.2 del mice and is likely to have multiple effects via engagement of downstream signaling pathways (Figure 1H).

**Figure 1.**
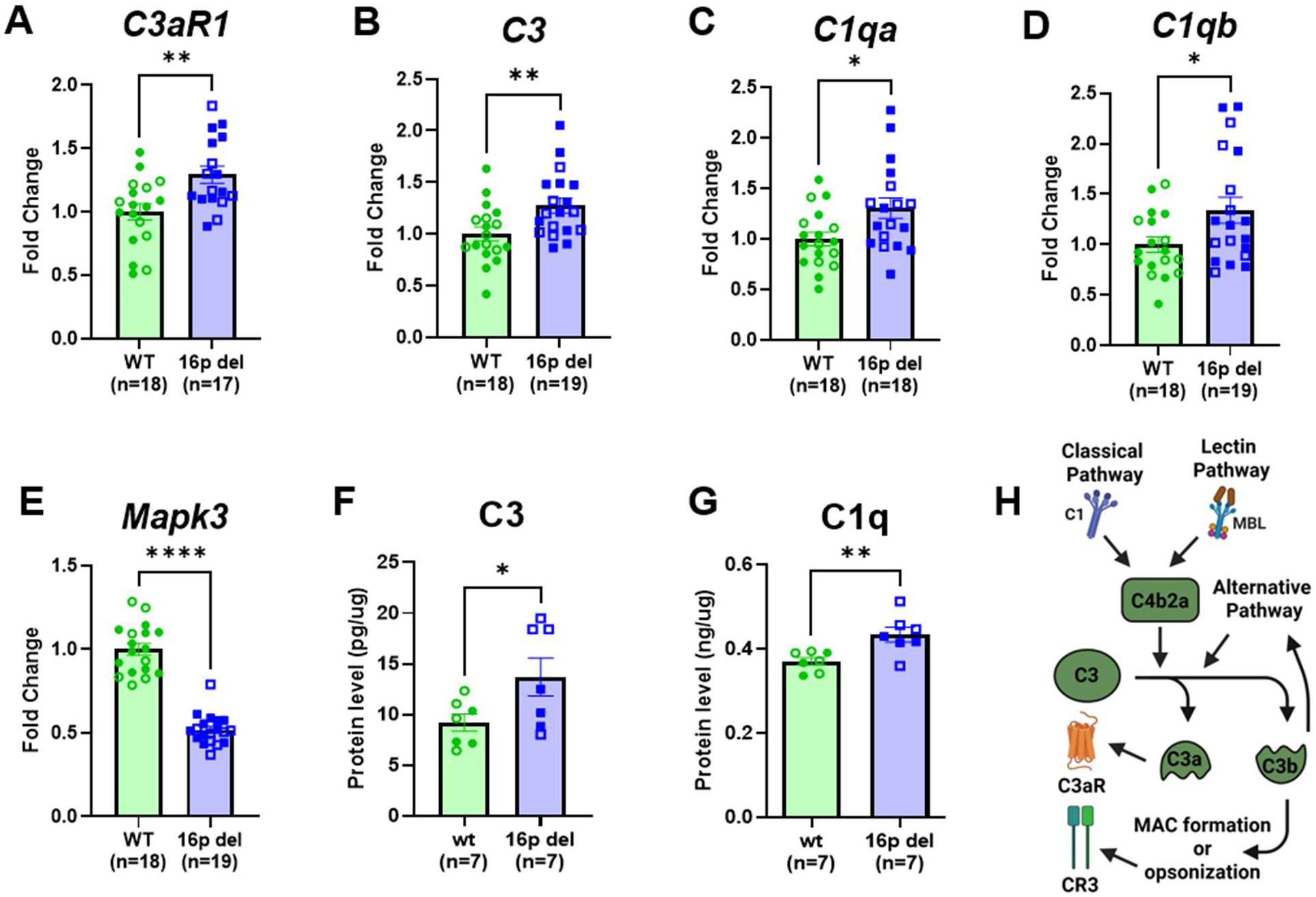
Complement expression is increased in the striatum of 16p11.2 del mice. A-D) Complement components *C3aR1*, *C3*, *C1qa*, and *C1qb* are upregulated in the striatum of 16p11.2 del mice at the mRNA level (*C3aR1*, t(33)=3.151, p=0.0034; *C3*, t(35)=2.780, p=0.0087; *C1qa*, t(34)=2.487, p=0.0180; *C1qb*, t(35)=2.280, p=0.0288). E) *Mapk3*, a gene within the 16p11.2 region, shows the expected 50% reduction in expression from WT levels (t(35)=11.96, p<0.0001). F,G) Complement components C3 and C1q are increased at the protein level in the striatum of 16p11.2 del mice (C3, t(12)=2.193, p=0.0487; C1q, t(12)=3.185, p=0.0078). H) A simple schematic representation of the complement cascade shows the convergence of the classical, lectin, and alternative pathways on proteolysis of C3 to mediate downstream signaling. Data is analyzed using student’s t-test (* indicates p < 0.05, ** indicates p < 0.01, **** indicates p < 0.0001). Error bars represent mean +/- standard error of the mean. Closed data points represent data obtained from male animals, and open data points indicate data obtained from female animals.

### 3.2 Complement drives hyperactivity in 16p11.2 del mice

To determine the impact of complement activation on behavior relevant to NDDs, we use the C3aR blocker, SB290157, and examine whether home cage hyperactivity found in both male and female 16p11.2 del mice is altered by C3aR antagonism (Figure 2A). Computational analysis predicts that SB290157 is permeable to the blood-brain barrier (Supplementary Figures 3A, B). Consistent with previous observations, 16p11.2 del mice are hyperactive in the home cage compared to WT littermates (Figures 2B, C). However, SB290157 treatment (10mg/kg, i.p.) reduces activity in 16p11.2 del animals (Figures 2D, E) whereas locomotor activity remains unchanged in WT animals (Figures 2F, G). The activity of 16p11.2 del animals in the presence of SB290157 is reduced to the level of WT mice (Figures 2H, I), showing that inhibition of an overactive complement system ameliorates hyperactive behavior in 16p11.2 del mice. Thus, complement activation drives NDD-relevant hyperactivity in 16p11.2 del mice.

**Figure 2.**
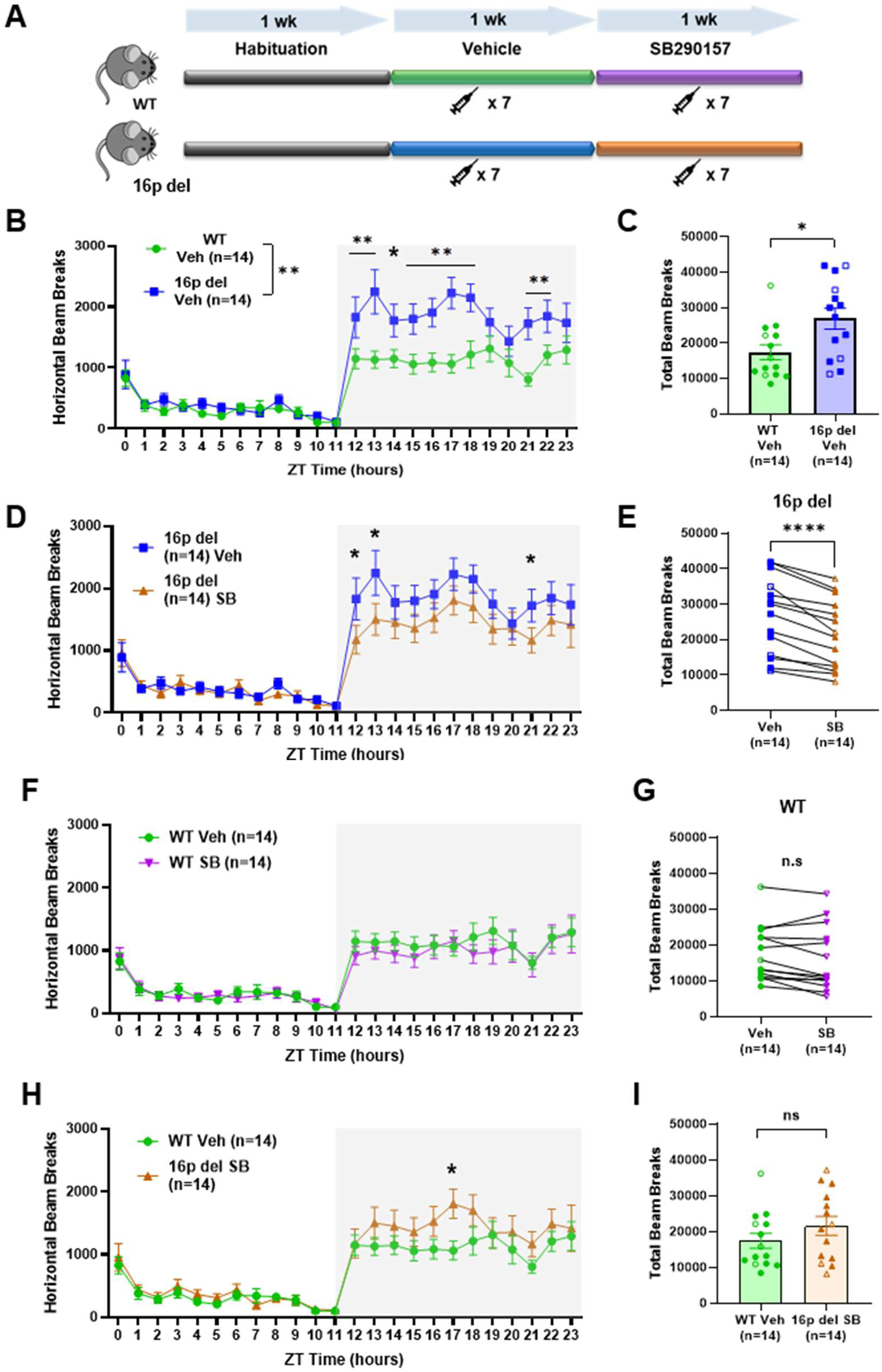
C3aR antagonist reduces hyperactivity in 16p11.2 del mice. The C3aR antagonist SB290157 (10 mg/kg, i.p.) is delivered at the onset of the dark cycle daily for one week to WT and 16p11.2 del animals (A). 16p11.2 del mice are hyperactive (B,C), and this hyperactivity is reduced upon antagonist administration (D,E) to WT levels (H,I). Activity in WT mice is not affected by SB290157 treatment (F,G). B) Main effect of genotype, F(1, 26)=6.983, p=0.0138; main effect of time, F(23, 598)=37.17, p<0.001; genotype x time interaction, F(23, 598)=3.997, p<0.001. Post hoc shows significant differences between vehicle injected WT and 16p11.2 del mice in the 12-18- and 21-22-hour time bins. C) 24-hour plot shows increased activity in 16p11.2 del mice (t(26)=2.643, p=0.0138). D) Main effect of genotype, F(1, 26)=1.789, p=0.1926; main effect of time, F(23, 598)=34.93, p<0.001; genotype x time interaction, F(23, 598)=1.186, p=0.2502. E) 24-hour plot with paired t-test shows that hyperactivity is reduced upon antagonist administration in 16p11.2 del mice (t(13)=6.727, p<0.0001). F) Main effect of treatment, F(1, 624)=2.341, p=0.1265; main effect of time, F(23, 624)=17.15, p<0.0001; treatment x time interaction, F(23, 624)=0.3142, p=0.9992. G) 24-hour plot with paired t-test shows no change in WT activity following SB290157 treatment (t(13)=2.045, p=0.0617). H) Main effect of genotype and treatment, F(1, 26)=1.601, p=0.2170; main effect of time, F(5.887, 153.1)=27.80, p<0.0001; genotype and treatment x time interaction, F(23, 598)=1.091, p=0.3498. Post hoc shows a significant difference between vehicle injected WT and SB290157 treated 16p11.2 del mice in the 17-hour time bin. I) 24-hour plot shows no difference in activity between vehicle injected WT and SB290157 treated 16p11.2 del mice (t(26)=1.265, p=0.2170). Two-way ANOVA with Fisher’s LSD *post-hoc* tests (B,D,F,H), student’s t-test (C,I), and paired t-test (E,G) are used for analyses (* indicates p < 0.05, ** indicates p < 0.01, **** indicates p < 0.0001). Error bars represent mean +/- standard error of the mean. Closed data points represent data obtained from male animals, and open data points indicate data obtained from female animals.

### 3.3 16p11.2 del produces a pro-inflammatory cytokine environment and striatal microglia activation

Because C3aR signaling is pro-inflammatory (30), we utilize an inflammatory cytokine array to determine the impact of 16p11.2 del on the inflammation state of the striatum (Supplementary Figure 4). This analysis reveals the upregulation of multiple cytokines, including IL-4, ILX, MCP-1, SDF-1, sTNF RI, and sTNF RII (Figures 3A-F, Supplementary Figure 4), suggesting that complement activation is part of a broader pro-inflammatory response to 16p11.2 del in the striatum. To further explore the impact of this inflammatory phenotype, we investigate microglia because, consistent with previous reports (31,32), C3aR is expressed predominantly in microglia (Supplementary Figure 5A). C3 expression, on the other hand, is localized primarily to astrocytes (Supplementary Figure 5C) rather than microglia (Supplementary Figure 5B). To assess the impact of 16p11.2 del on microglia, we perform morphological analyses of striatal microglia using the MicrogliaMorphology analysis package on Iba1 stained microglia (29). This analysis categorizes microglia into four morphological states indicative of their level of activation (Figure 4A, Supplemental Figures 6A-D). We found that microglia in the 16p11.2 del striatum are more likely to be amoeboid, the morphology most indicative of microglial activation (Figure 4B). This increase in amoeboid microglia is accompanied by a decrease in hypertrophic microglia, the second most active morphology, and no change in the percentage of rod-like or ramified microglia (Figures 4C-E). Furthermore, microglia in the 16p11.2 del striatum exhibit a trend, though not statistically significant, toward increased density (Supplemental Figure 6E) and Iba1 signal intensity (Supplemental Figure 6F), both phenotypes indicative of microglial activation. Together, these data suggest that 16p11.2 del has a subtle effect on striatal microglia activation where some microglia that are prone to activation become further activated.

**Figure 3.**
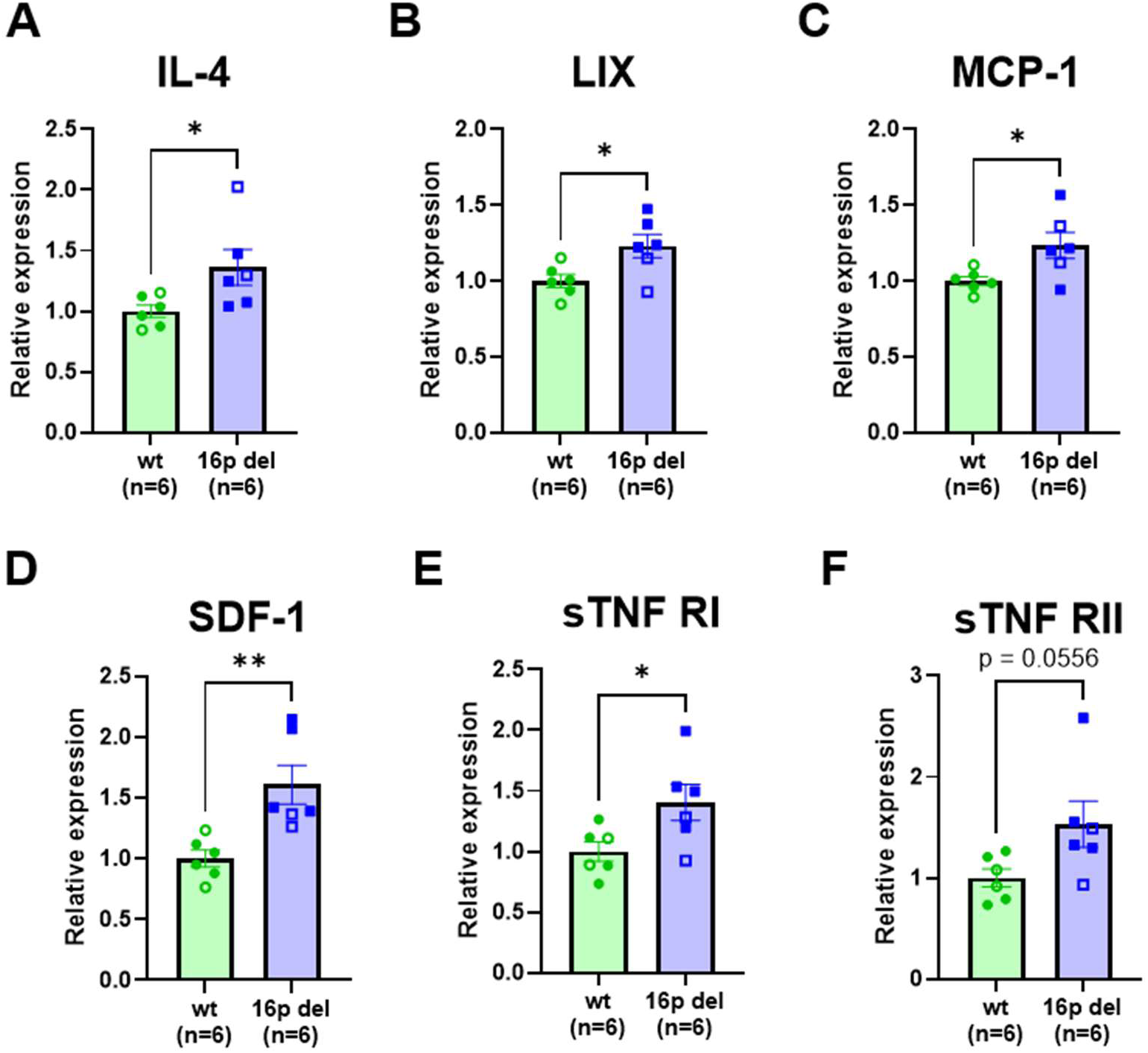
Inflammatory factors are upregulated in the striatum of 16p11.2 del mice. A-F) Levels of cytokines IL-4, LIX, MCP-1, SDF-1, sTNF RI, and sTNF RII are increased in the striatum of 16p11.2 del mice (IL-4, t(10)=2.299, p=0.0443; LIX, t(10)=2.613, p=0.0259; MCP-1, t(10)=2.552, p=0.0288; SDF-1, t(10)=3.510, p=0.0056; sTNF RI, t(10)=2.407, p=0.0369; sTNF RII, t(10)=2.166, p=0.0556;). Data is analyzed using student’s t-test (* indicates p < 0.05, ** indicates p < 0.01). Error bars represent mean +/- standard error of the mean. Closed data points represent data obtained from male animals, and open data points indicate data obtained from female animals.

**Figure 4.**
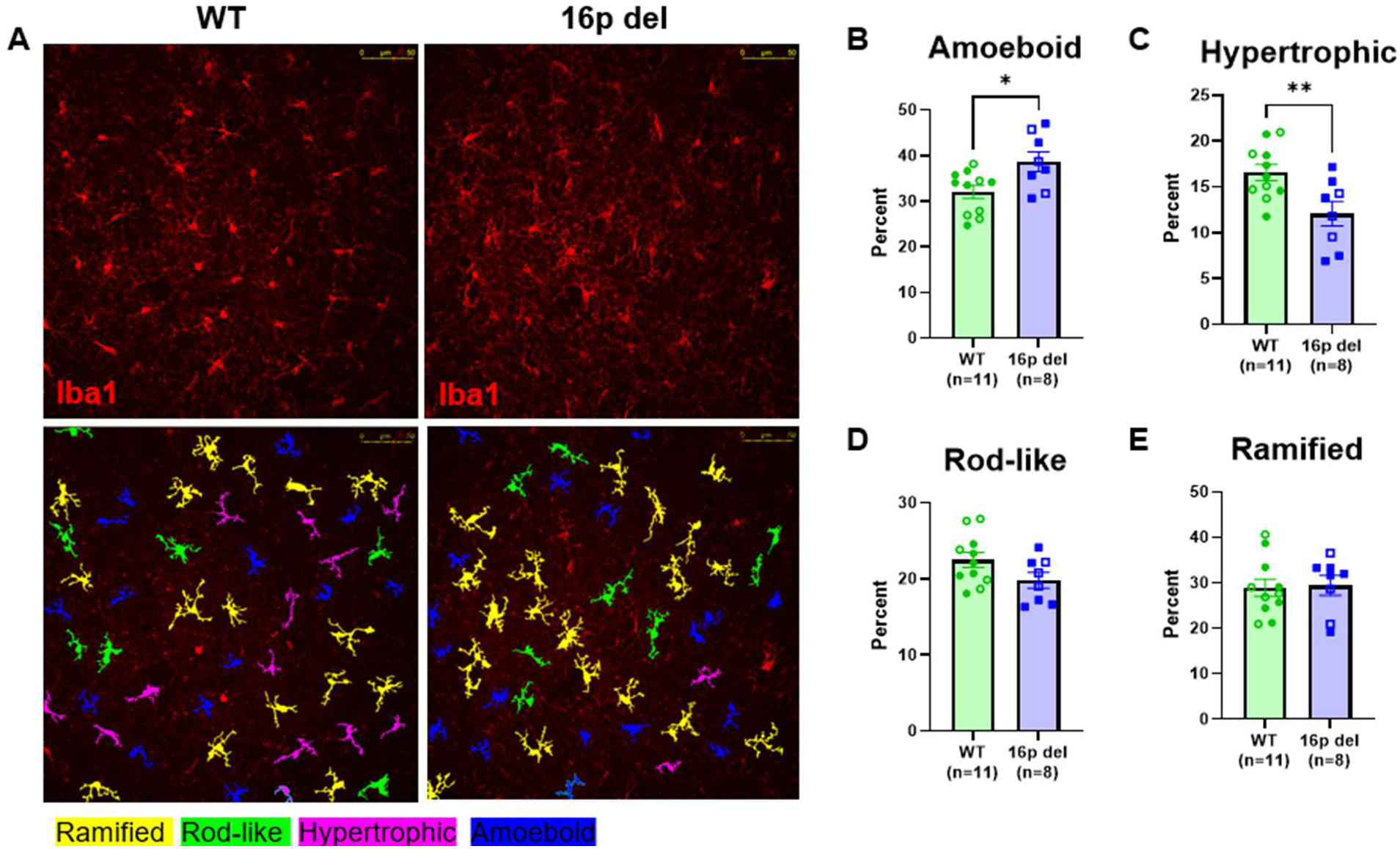
Microglia within the 16p11.2 del striatum have altered morphology. A) Representative images (20X magnification) of striatal microglia morphology that are profiled in WT and 16p11.2 del mice using the MicrogliaMorphology ImageJ macro and R package, which assigns individual microglia to one of four morphologies: ramified, rod-like, hypertrophic, or amoeboid. B,C) 16p11.2 del animals have an increased percentage of amoeboid microglia along with a decrease in hypertrophic cells (amoeboid, t(17)=2.673, p=0.0160; hypertrophic, t(17)=2.920, p=0.0095). D,E) The percentage of rod-like and ramified microglia are unchanged (rod-like, t(17)=1.809, p=0.0882; ramified, t(17)=0.1851, p=0.8553). Data is analyzed using student’s t-test (* indicates p < 0.05, ** indicates p < 0.01). Error bars represent mean +/- standard error of the mean. Closed data points represent data obtained from male animals, and open data points indicate data obtained from female animals.

### 3.4 Inhibiting microglial activation does not impact locomotion of 16p11.2 del mice

To assess the behavioral impact of microglial activation in 16p11.2 del animals, we inhibit pro-inflammatory microglial activation pharmacologically using minocycline and observe the effect on activity in the home cage (Figure 5A). 16p11.2 del mice treated with minocycline (50 mg/kg, i.p.) remain hyperactive compared to WT mice (Figures 5B, C). Minocycline treatment does reduce activity in 16p11.2 del mice, particularly at the onset of the dark cycle when mice are most active (Figures 5D, E). However, minocycline also produces a similar effect in WT mice (Figures 5F, G). These data suggest that hyperactivity in 16p11.2 del mice does not depend on canonical pro-inflammatory activation of microglia. Taken together with the effect of pharmacological inhibition of C3aR on hyperactive behavior, our data suggests that specific mechanisms downstream of complement signaling lead to NDD-relevant hyperactive behavior in 16p11.2 del mice, potentially through non-canonical activation of microglia, though the involvement of other cell types remains possible.

**Figure 5.**
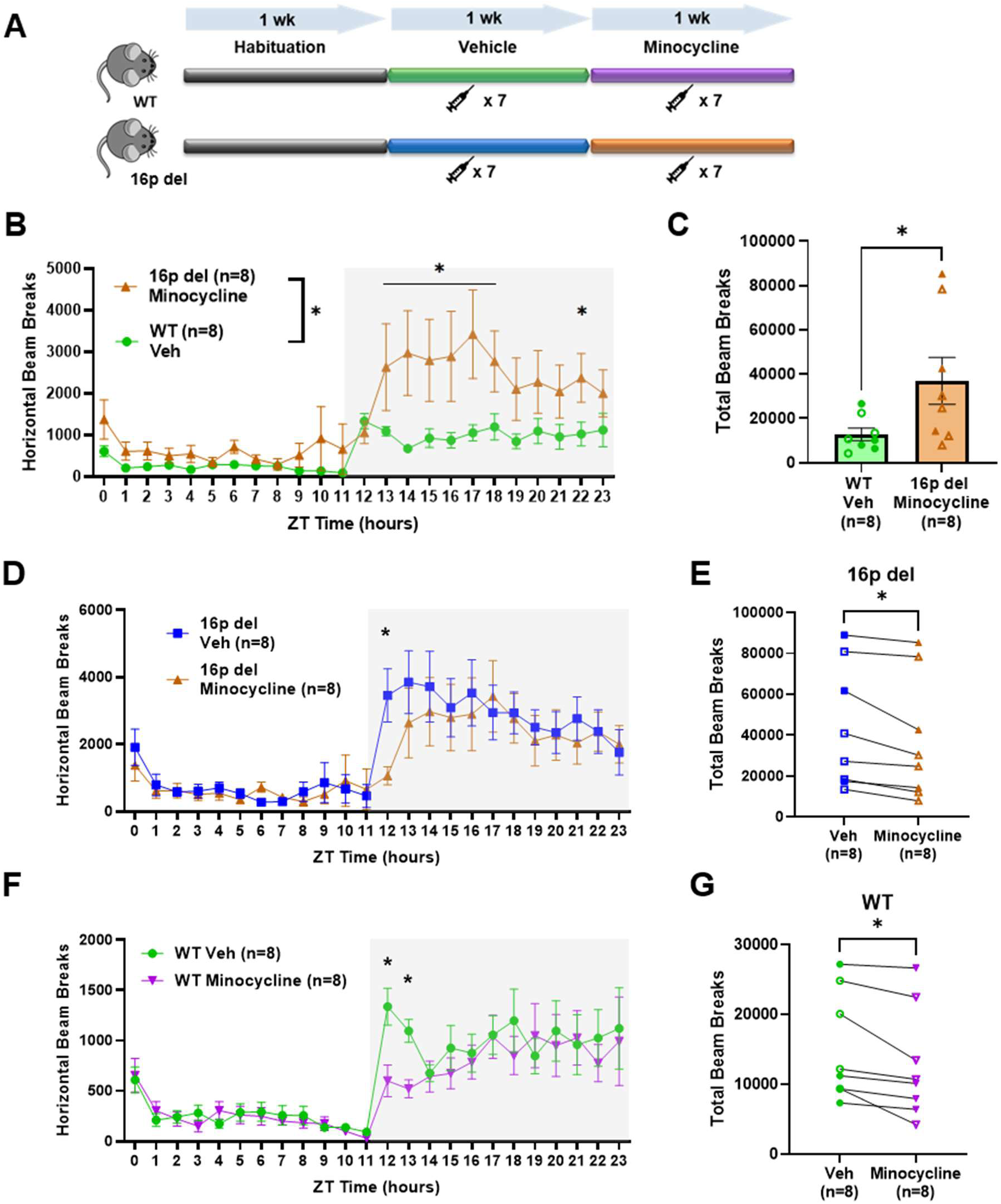
16p11.2 del mice remain hyperactive after minocycline treatment. Minocycline (50 mg/kg, i.p.) is given chronically to WT and 16p11.2 del mice to inhibit pro-inflammatory activation of microglia (A). Minocycline treatment does not lower 16p11.2 del activity to WT level (B,C). Reduced activity is observed in both 16p11.2 del (D,E) and WT (F,G) mice treated with minocycline particularly at the onset of the dark cycle. B) Main effect of genotype and treatment, F(1, 336)=44.90, p<0.0001; main effect of time, F(23, 336)=4.467, p<0.0001; genotype and treatment x time interaction, F(23, 336)=1.277, p=0.1796. Post hoc analyses show a significant difference between vehicle injected WT and minocycline treated 16p11.2 del mice in the 13-18- and 22-hour time bins. C) 24-hour plot shows significantly higher activity in minocycline treated 16p11.2 del mice compared to vehicle injected WT mice (t(14)=2.212, p=0.0441). D) Main effect of treatment, F(1, 336)=2.422, p=0.1206; main effect of time, F(23, 336)=6.542, p<0.0001; treatment x time interaction, F(23, 336)=0.4737, p=0.9826. Post hoc analyses show a significant decrease in activity following minocycline treatment in 16p11.2 del mice in the 12-hour time bin. E) 24-hour plot shows that activity is reduced upon minocycline administration in 16p11.2 del mice (t(7)=3.321, p=0.0127). F) Main effect of treatment, F(1, 336)=3.841, p=0.0508; main effect of time, F(23, 336)=8.365, p<0.0001; treatment x time interaction, F(23, 336)=0.6833, p=0.8623. Post hoc shows a significant decrease in activity following minocycline treatment in WT mice in the 12–13- hour time bins. G) 24-hour plot with paired t-test shows that activity was reduced upon minocycline administration in 16p11.2 del mice (t(7)=3.120, p=0.0168). Two-way ANOVA with Fisher’s LSD *post-hoc* test (B,D,F), student’s t-test (C), and paired t-test (E,G) are used for analyses (* indicates p < 0.05). Error bars represent mean +/- standard error of the mean. Closed data points represent data obtained from male animals, and open data points indicate data obtained from female animals.

## 4. Discussion

We demonstrate that the inhibition of an overactive complement system reduces hyperactive behavior in the 16p11.2 del mouse model. Although complement has previously been shown to be upregulated in both humans with ASD and relevant preclinical models (19,33,34), this is the first time that an overactive complement cascade has been shown to cause NDD-relevant behavior. These data open the door for new therapeutic strategies to ameliorate behavioral challenges associated with NDDs. They also highlight the 16p11.2 del model as a promising tool to understand the impacts of excess complement in the context of NDDs.

The cellular and molecular mechanisms downstream of C3aR in the 16p11.2 del model require further investigation. Microglia are responsible for the vast majority of C3aR expression in the brain (31,32), and they are also the primary cellular source of C1q in the CNS (35,36). C3aR activation in the CNS is most strongly associated with microglial inflammation downstream of G_i/o_ signaling (30,37,38). Our results indicate that striatal 16p11.2 del microglia exhibit a morphological signature of pro-inflammatory activation. However, minocycline treatment, which prevents microglial inflammation, does not reproduce the activity lowering effect of C3aR inhibition. This raises the possibility that C3aR signaling in other cell types contributes to hyperactivity in 16p11.2 del animals. Indeed, C3aR signaling can impact neurons as well as endothelial cells, particularly under pro-inflammatory conditions (39,40). Furthermore, the proximal cause of complement upregulation in the 16p11.2 del model is not known. Complement expression is controlled by multiple factors including ERK signaling and NFĸB transcription factor activity (41–43). Prior work has described ERK overactivation in 16p11.2 del mice, which is paradoxical, given that the gene coding for ERK1, *mapk3*, is within the 16p11.2 region (44–46). However, the role of ERK signaling in controlling complement expression in 16p11.2 del remains to be determined. Further studies will be required to pinpoint the cellular and molecular underpinnings of complement dysregulation in the 16p11.2 del model.

The impact of complement upregulation on the neural circuitry mediating hyperactivity in 16p11.2 del mice is unknown. We reported selective complement upregulation in the striatum, a region known to be impacted in both humans with 16p11.2 del and preclinical models (6,23,25,46–49). Our group recently demonstrated that hyperactivity specifically in male 16p11.2 del mice is mediated by the striatal indirect pathway made up of dopamine receptor D2 expressing medium spiny neurons (D2 MSNs), suggesting that the same behavioral phenotype in a single genetic model may be mediated by different underlying circuitry (25). Complement has been shown to selectively sculpt cortico-striatal circuits through microglia-mediated synaptic pruning in an HD model (26). Although more work is needed to establish that microglia-mediated synaptic pruning plays a role in the hyperactivity of 16p11.2 del mice, 16p11.2 del is known to impact synaptic pruning. Specifically, increased expression of CD47, the “don’t eat me” signal that counteracts complement mediated phagocytosis, has been observed in both induced pluripotent stem cells modeling 16p11.2 del and the prefrontal cortex of 16p11.2 del mice, inhibiting synaptic pruning and contributing to brain overgrowth and NDD-relevant behavior (50,51). This seems to contradict our observation of increased complement expression in the striatum that predicts increased pruning of cortico-striatal synapses and contributes to decreased D2 MSN activity, although these differences may be explained by the differential impact of 16p11.2 del on different cell types and in different brain regions. This model would not explain the male-specific contribution of D2 MSNs to hyperactivity in 16p11.2 del mice. However, previous work has shown that complement-mediated synaptic pruning has sex-specific impacts on striatal circuitry during strict developmental windows (52). Indeed, the action of sex hormones on microglia during development can have lasting impacts on neural circuits and behavior (53). More work is required to delineate the developmental and sex-specific impacts of complement upregulation on cortico-striatal circuits in the 16p11.2 del model and to determine how these processes may mediate hyperactivity and other NDD-relevant behavior.

Our results designate the complement system as a promising target for the development of new treatments for individuals with 16p11.2 del and other neurodevelopmental challenges. Although the use of SB290157 has been limited to research applications due to reported off-target effects (54), newer generations of C3aR antagonists with greater specificity are promising agents for preclinical development (55,56). Furthermore, ANX005, a therapeutic monoclonal antibody targeting C1q, has shown promising results in clinical trials for Guillain-Barré Syndrome (57). Additional preclinical studies are necessary to determine the potential of these strategies in individuals with 16p11.2 del, and more mechanistic studies will serve to both expand and refine the molecular targets amenable to therapeutic development. Our work opens the door for the advancement of agents targeting the complement system to treat the challenges associated with NDDs.

## Acknowledgements

Support for this work was provided by the National Institutes of Health (NIH F31 MH134542, NIH T32 GM067795, NIH R01 MH087463, NIH R01 DA056113) and the University of Iowa Hawkeye Intellectual and Developmental Disabilities Research Center (HAWK-IDDRC, NIH P50 HD103556). TA is supported by the Roy J. Carver Directorship of the Iowa Neuroscience Institute. We thank Drs. Annie Vogel Ciernia and Jennifer Kim for their guidance in analyzing microglia using their MicrogliaMorphology package. We would also like to acknowledge the Neural Circuits and Behavior Core in the Iowa Neuroscience Institute, which provided equipment and facilities to support behavioral and imaging experiments. Figures created with BioRender.com.

## CRediT author statement

**Benjamin Kelvington:** Conceptualization, Formal analysis, Investigation, Writing - Original Draft, Funding acquisition; **Jaekyoon Kim:** Conceptualization, Formal analysis, Investigation, Writing - Review & Editing; **Regan Fair:** Formal analysis, Investigation; **Marie Gaine:** Conceptualization, Writing - Review & Editing; **Ted Abel:** Conceptualization, Writing - Review & Editing, Supervision, Funding acquisition.

