## Supplemental Material for "Complement contributes to hyperactive behavior in the 16p11.2 hemideletion mouse model"

### Supplementary Figures

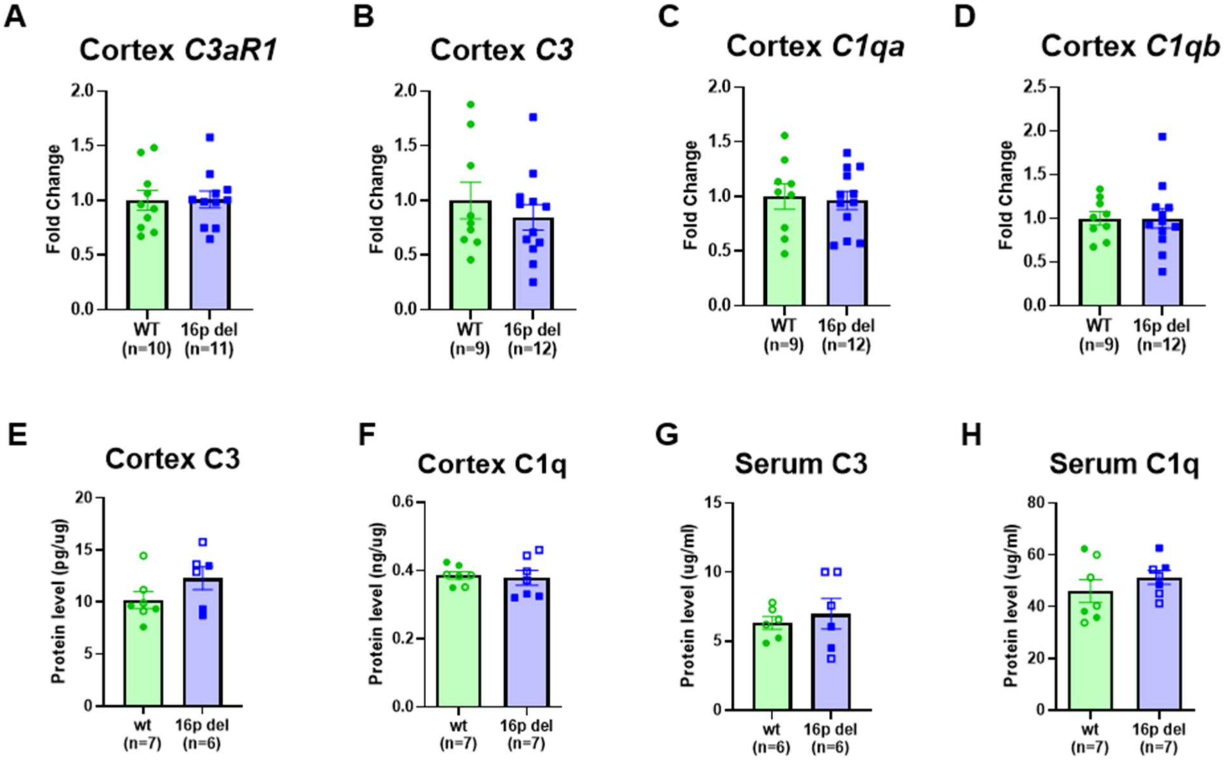


**Supplementary Figure 1. Complement upregulation is localized to striatum of 16p11.2 del mice.**

A-D) mRNA levels of complement components *C3aR1*, *C3*, *C1qa*, and *C1qb* are unchanged in the cortex of 16p11.2 del mice compared to WT (*C3aR1*, t(19)=0.08149, p=0.9359; *C3*, t(19)=0.7807, p=0.4446; *C1qa*, t(19)=0.2569, p=0.8000; *C1qb*, t(19)=0.01090, p=0.9914). Complement proteins C3 and C1q are also observed at WT levels in both the cortex (E,F) and serum (G,H) of 16p11.2 del mice (C3 cortex, t(11)=1.559, p=0.1473; C1q cortex, t(12)=0.2988, p=0.7702; C3 serum, t(10)=0.5502, p=0.5943; C1q serum, t(12)=1.015, p=0.3300). Data is analyzed using student's t-test. Error bars represent mean +/- standard error of the mean. Closed data points represent data obtained from male animals, and open data points indicate data obtained from female animals.

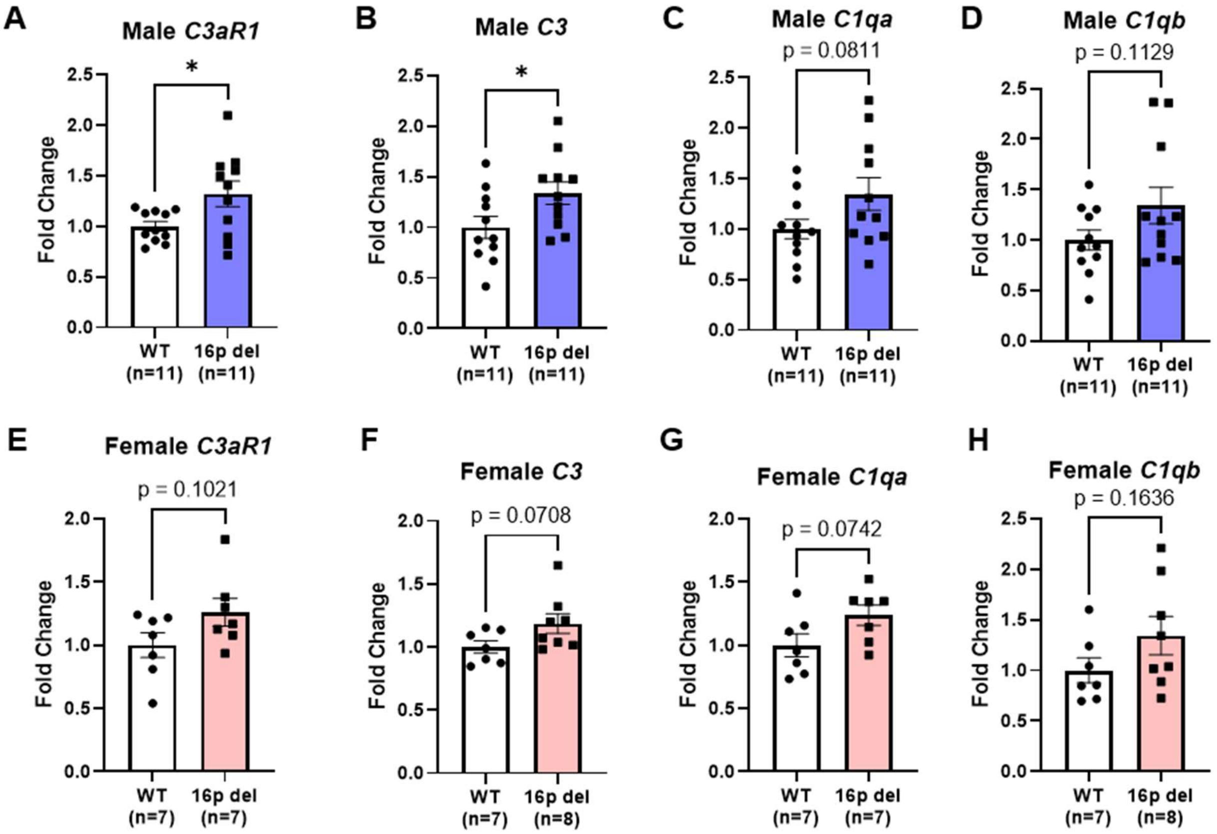


**Supplementary Figure 2. Male and female 16p11.2 del mice express similar levels of striatal complement.**

Expression of complement component mRNA is elevated in both male (A-D) and female (E-H) 16p11.2 del animals compared to WT (male *C3aR1*, t(20)=2.398, p=0.0264; male

*C3*, t(20)=2.197, p=0.0400; male *C1qa*, t(20)=1.837, p=0.0811; male *C1qb*, t(20)=1.658, p=0.1129; female *C3aR1*, t(12)=1.770, p=0.1021; female *C3*, t(13)=1.968, p=0.0708; female *C1qa*, t(12)=1.956, p=0.0742; female *C1qb*, t(13)=1.477, p=0.1636). Data is analyzed using student's t-test (* indicates $p < 0.05$). Error bars represent mean +/- standard error of the mean.

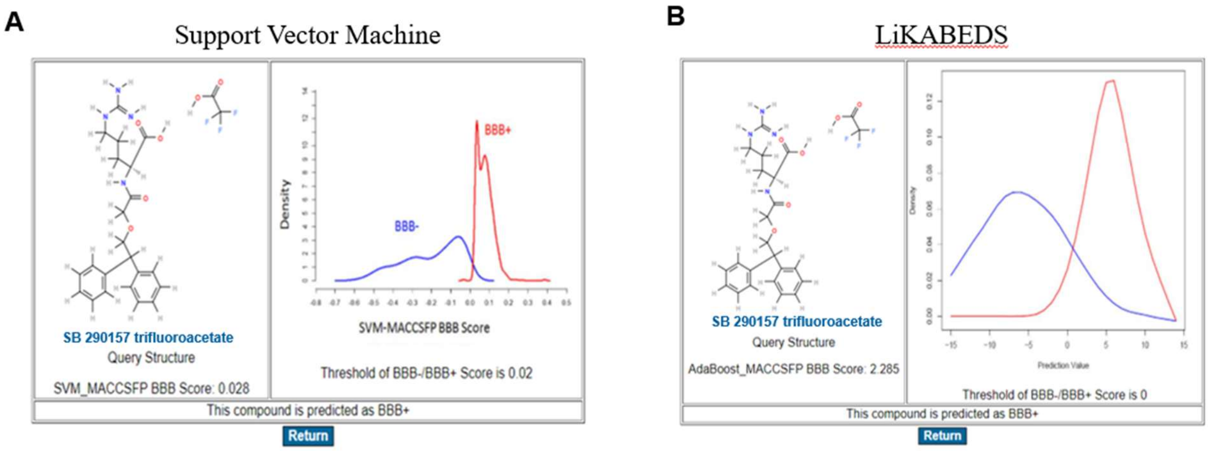


**Supplementary Figure 3. C3aR antagonist SB290157 is predicted to penetrate the blood-brain barrier.**

The likelihood of SB290157 penetrating the blood brain barrier (BBB) is calculated using the online BBB server (http://www.cbligand.org/BBB) version of (A) the Support Vector Machine (SVM_MACCSFP BBB Score: 0.028) and (B) LiKABEDS (AdaBoost_MACCSFP BBB Score: 2.285) algorithms developed by Liu et al (1). These algorithms are based on four types of fingerprints for 1593 training compounds. In both cases, BBB penetration is predicted to occur.

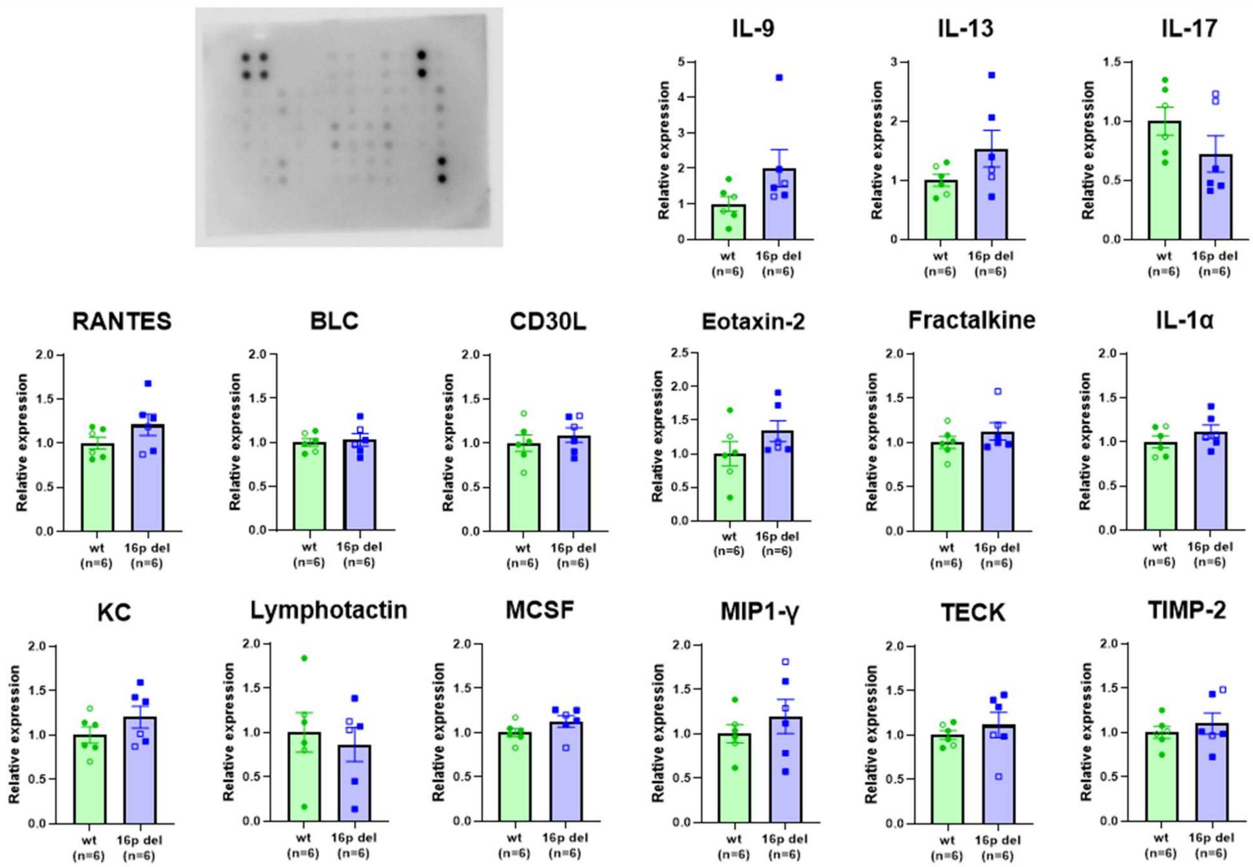


**Supplementary Figure 4. Inflammatory environment of 16p11.2 del striatum profiled by cytokine array.**

Expression of inflammatory cytokines is assessed using a cytokine array (IL-9, $t(10)=1.790$, $p=0.1038$; IL-13, $t(10)=1.640$, $p=0.1320$; IL-17, $t(10)=1.427$, $p=0.1841$; RANTES, $t(10)=1.490$, $p=0.1670$; BLC, $t(10)=0.3551$, $p=0.7299$; CD30L, $t(10)=0.7311$, $p=0.4815$; Eotaxin-2, $t(10)=1.424$, $p=0.1850$; Fractalkine, $t(10)=1.064$, $p=0.3125$; IL-1α, $t(10)=1.126$, $p=0.2864$; KC, $t(10)=1.323$, $p=0.2153$; Lymphotactin, $t(10)=0.4579$, $p=0.6568$; MCSF, $t(10)=1.591$, $p=0.1427$; MIP1-y, $t(10)=0.8861$, $p=0.3964$; TECK, $t(10)=0.7545$, $p=0.4680$; TIMP-2, $t(10)=0.7149$, $p=0.4910$). Data is analyzed using student's t-test. Error bars represent mean +/- standard error of the mean. Closed data points represent data obtained from male animals, and open data points indicate data obtained from female animals.

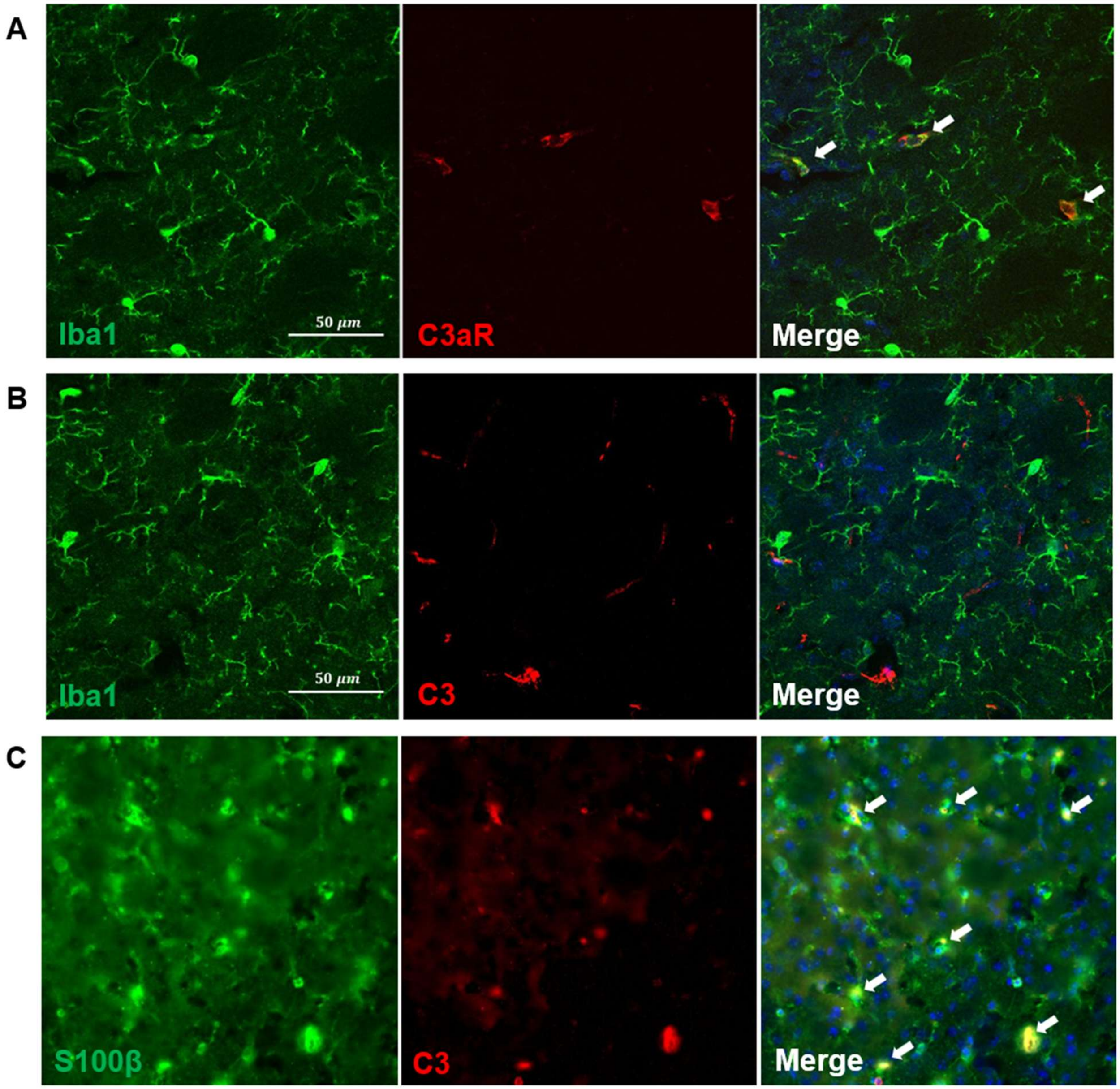


**Supplementary Figure 5. Cellular localization of complement pathway factors in the striatum.**

Immunohistochemistry is used to assess the cellular localization of complement proteins in the striatum. Representative images are shown at 40X magnification. C3aR co-localizes with the microglia marker Iba1 (A). C3 does not co-localize with Iba1 (B) but does co-localize with the astrocyte marker S100β (C).

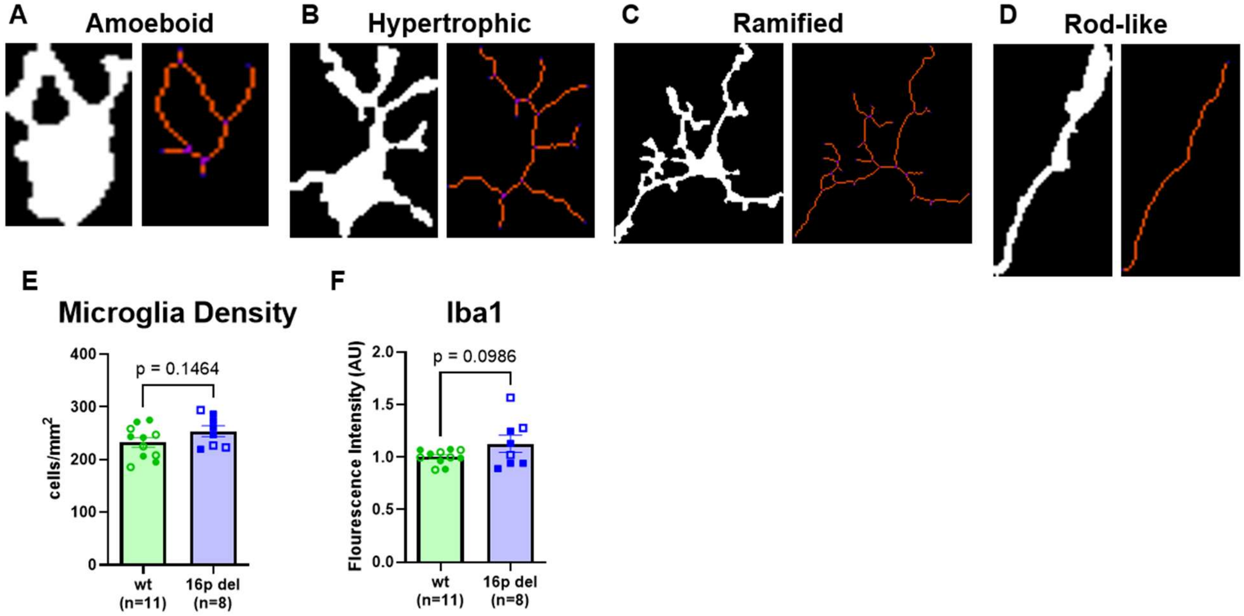


**Supplementary Figure 6. Microglia morphology analysis profiles microglia in 16p11.2 del striatum.**

MicrogliaMorphology ImageJ macro and R package assigns individual microglia to one of four morphologies: ramified, rod-like, hypertrophic, or amoeboid (A-D). Analysis of Iba1 labeled microglia reveals trends toward an increase in microglia density (E, t(17)=0.1464, p=0.1464) and Iba1 signal intensity (F, t(17)=1.747, p=0.0986). Data is analyzed using student's t-test. Error bars represent mean +/- standard error of the mean. Closed data points represent data obtained from male animals, and open data points indicate data obtained from female animals.

**Supplementary Table 1. qPCR Primer sequences**

| | | |
|---|---|---|
| *C3ar1* | Forward | TCG ATG CTG ACA CCA ATT CAA |
| | Reverse | TCC CAA TAG ACA AGT GAG ACC AA |
| *C3* | Forward | GCG TAA TAC GAC TCA CTA TAG GGC AGG GGA GTA TAT TGA AGC CAG |

| | | |
|---|---|---|
| | Reverse | GCG ATT TAG GTGACA CTA TAG TCT ATC TAC TCC AGA GGC CAG C |
| *C1qa* | Forward | AAA GGC AAT CCA GGC AAT ATC |
| | Reverse | TGG TTC TGG TAT GGA CTC TCC |
| *C1qb* | Forward | CGT CGG CCC TAA GGG TAC T |
| | Reverse | GGG GCT GTT GAT GGT CCT C |
| *Mapk3* | Forward | CCA AAG CTC TTG ACC TGC TG |
| | Reverse | CCA GCG CTT CCT CTA CTG TG |
| *Tubulin* | Forward | ATG CGC GAG TGC ATT TCA |
| | Reverse | CAC CAA TGG TCT TAT CGC TGG |
| *Gapdh* | Forward | TGC ACC ACC AAC TGC TTA GC |
| | Reverse | GGC ATG GAC TGT GGT CAT GA |
